# Computational mutagenesis by using AUTO-MUTE 2.0 to examine X-ray crystallography structures for p53 and the impact of its resolution on predictions

**DOI:** 10.1101/2023.01.03.522606

**Authors:** Shaimaa Sait, Isoif Vaisman

**Affiliations:** School of system biology, George Mason University, Manassas, Virginia

## Abstract

The mutation of P53 is found in 50% of all human cancers, affecting the protein structure conformation, which further impacts its function. Identifying the appropriate protein structure for future research has been challenging. Many X-ray crystallography structures exist in the Protein Data Bank (PDB) for a given protein. We compared two proteins with different resolutions to determine whether the differences between the two structures were statistically significant. The present study used mutagenesis to study two structures of the protein P53 with various resolutions to assess how the conformation of the protein changes upon mutation. We used an AUTO-MUTE 2.0, which is a computational geometry method based on the Delaunay tessellation for predicting protein structures. The current study compares the prediction results obtained from different predictive methods. We used logistic regression models and corresponding receiver operating characteristic (ROC) curves to model the differences between structures using the variables available at PDB. We conclude from this study that x-ray crystallography is a sensitive and advanced technique, and the resulting proteins at different resolutions are similar.

## INTRODUCTION

Mutations in the gene P53 (also known as the guardian of the genome) are considered the most common cause of human cancer. Approximately 50% of cancers in humans have P53 mutations (Baugh et al. 2016, 2018). P53 plays a crucial role in the proliferation and apoptosis of cells. A mutation of this protein will profoundly affect cell function and behavior. Substitutions or deletions of the nucleotide can result in the loss of function. P53 prevents tumor development by regulating cell-cycle arrest and apoptosis (Kim and Lozano 2018). Many mutations in P53 are in the DNA binding domain. Under stressful situations like DNA damage, the wild protein increases in the nucleus and regulates genes involved in the cell cycle’s checkpoints. Thus, P53 is recognized as the central player in cancer by preventing tumors and repairing DNA damage. DNA damage cannot be repaired when P53 is inactive. Therefore, the active version of P53 will prevent cancer (Armstrong 2015; Muller and Vousden 2013).

The practice of in silico mutagenesis has increasingly been observed in recent years. Using SDM, HoTMuSiC, I-Mutant, SDM, and iStable2 methods, one can predict the effect of amino acid replacement of single residues most reliably and rapidly (Baugh et al. 2016). This study aimed to determine the impact of single-point mutations on the overall structure and function of the DNA domain for protein p53 by analyzing its structure for different resolutions.

It is challenging to determine the appropriate structure among tens of the protein structures available in the PDB database. The study identifies two protein structures (1TSR and 4IBY) (Cho et al. 1994; Eldar et al. 2013) that vary in resolution, which can be useful for studying comprehensive mutagenesis to determine the effect of differences in protein structures on functional predictions of protein mutants. The study employs a method combining structure-based features as in Delaunay tessellation with training statistical regression models, which are present in AUTO-MUTE 2.0, to confirm for predicting the functional change of proteins upon single residue replacement. AUTO-MUTE 2.0 has developed different predictors models based on energy Stability Change (SC), Thermostability (TC), Activity Changes (AC), and Disease Potential (DP) (Masso and Vaisman 2010a). The models were developed using the Java-based Weka software package to implement classification and regression statistical machine learning algorithms. This study aims to examine whether protein resolution influences the interpretation of protein structure. The AUTO-MUTE 2.0 free download is available on (http://proteins.gmu.edu/automute). AUTO-MUTE 2.0 is a free software application that relies primarily on computational geometry and statistical machine learning tools such as Q-hull (http://www.qhull.org/) (Barber, Dobkin, and Huhdanpaa 1996) and Weka (http://www.cs.waikato.ac.nz/ml/weka/) (Frank et al. 2004). These tools are not platform-independent and are used without modification by the software (Masso and Vaisman 2014).

## METHOD

This method focuses on the replacement of a single residue on the local structural effects. It uses the Qhull program to identify all residue positions that are structural neighbors (adjacent) to the residue undergoing replacement. A unique feature vector that describes a protein mutation is composed of individual component values corresponding to the mutation and its six closest neighbors, including critical measures quantifying structural effects. For in silico mutagenesis, a four-body energy function is used to determine structural perturbations. The predictive models incorporate the feature vectors of the single residue mutations as inputs into statistical machine-learning algorithms implemented in Weka software (Masso and Vaisman 2014).

The AUTO-MUTE 2.0 program converts a file containing user-supplied requests for single residue substitution into a numerical file containing their respective feature vectors. First, the Qhull method identifies the six closest structural neighbors of each position in each protein structure undergoing a mutation. Following this, the support programs prepared were used to determine the attribute values for each residue mutation prediction request. Next, a table is generated with predictions made by AUTO-MUTE 2.0 based on the file that contains feature vectors for requested mutations. AUTO-MUTE 2.0 programs use predictive models trained on thousands of single residue mutations and experimentally demonstrate their functional consequences (stability change, activity change, or human nsSNP disease potential). Finally, these trained Weka models predict the functional effects of single residue mutations (Masso and Vaisman 2010b).

The PDB files are what AUTO-MUTE 2.0 uses as its input format in conjunction with mutant files. PDB files contain atomic coordinates and information regarding protein structure. For each amino acid, the α-carbon coordinates are located on the first carbon atom that attaches to a functional group. An AUTO-MUTE 2.0 algorithm is applied to determine which amino acid represents each component. The PDB file describes how the α-carbon is constrained to points in three-dimensional space. In order to classify the quadruplets of each amino acid in the protein, the Delaunay tessellation is applied, where each point is represented as a vertex. The tetrahedral of six edges smaller than 12 Å is used as quadruplets.

Let (i, j, k, and l) quadruplet of each residue; the Delaunay tessellation algorithm is used to measure the frequency of occurrence (f_ijkl_) for approximately 8855 quadruplets formed from the naturally occurring twenty amino acids by utilizing the training samples of more than 1400 high-resolution structures with short length sequence and similarity obtained from PDB. For the formation of quadruplets, multinominal distribution (n=4) is used to measure the excepted rate of frequency p(ijkl) (Masso and Vaisman 2010a). Finally, a log-likelihood score calculates the tendency for each quadruplet form, calculated by equation 1. Applying the inverted Boltzmann principle. Score (S) is the proportional interaction energy for the quadruplet of residues. 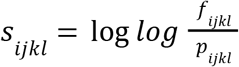. where ∑_*n*=1_ ^20^ a _*n*_ = 1 and ∑_*n*=1_ ^20^ *t _n_* = 4. Here, a & n represents the proportion of all residues comprising the 1417 proteins that are of type n, and *t _n_* is the number of occurrences of residue type n in the (i, j, k, and l) quadruplet

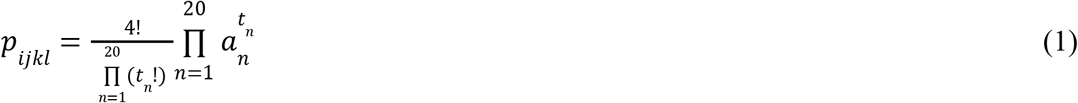

Each amino acid’s residual environment (RE) score is determined in a protein by combining the quadruplets’ log-likelihood score, which formed the vertex and produced the 3D-ID potential profile (Masso and Vaisman 2010b). Computational mutagenesis is modified to the residue mark at a particular place of interest and the residual environment score. Only the mutated and neighboring amino acid residues are observed to alter their environment score of 3D-ID potential profile (Bowie, Lüthy, and Eisenberg 1991). The differences in vectors of nonmutated and mutated residues of protein as a residual profile and its components of environment changes (EC) score count residues’ perturbations in protein (Pettersen et al. 2004).

The Delaunay tessellation is based on a classical computational geometry approach that presents the quadruplets of 3D nearest neighbor residues. The predictive models for stability change are based on Combining an energy approach to machine learning to develop the accuracy of Gibbs Free Energy Changes (ΔΔG) and melting temperature changes (ΔTm). The statistical machine learning algorithms are applied using the Weka software package (Frank et al. 2004). The initial stability modification database ΔTm included 1791 single amino acid substitutions in 68 different structural proteins derived from the PDB. For predicting the ΔΔG, trained random forest (RF) and support vector machine (SVM) classifiers were introduced adaptively boosted C4.5 decision tree and SVM classifiers for ΔTm. Tree regression (REPTree) and support vector regression SVR models, on the other hand, have been used on all three servers to forecast absolute stability transition values (Masso and Vaisman 2010b).

On the other hand, the activity changes of the target protein due to a single mutation are predicted using activity changes algorithms in the AUTO-MUTE 2.0 webserver. This server was predicted to change the stability of protein due to mutation in the protein sequence. The prediction shows that either single amino acid replacement at a specific position could affect or unaffected the structure of the target protein. The Random Forest follows this server as a classified model to evaluate stability changes (Cheng, Randall, and Baldi 2006). Human nsSNP disease potential is an automated server envisaging a human nucleotide polymorphism (nsSNP) in a coding area that induces an amino acid replacement causes or is only a neutral disease (Apweiler et al. 2004). The approach builds on the intuitive notion that nsSNPs, which are benign or contribute to abnormal protein production and disease, are related to comparative structural variations from the wild. The Random Forest (RF) algorithm is used for a monitored classification model on the server (Barenboim et al. 2008).

Measuring performance is crucial to Machine Learning. ROC (Receiver Operating Characteristics) curves and AUC (Area Under the Curve) are used to visualize the performance of multi-class classification problems (Niroula and Vihinen 2016). The ROC is a probability curve, while the AUC describes the degree or measure of separability. AUC indicates how well the model can distinguish between classes. A model’s sensitivity is defined as its ability to identify a structure that belongs to the stable within energy changes, stable within thermostability, unaffected activity, and neutral to disease. Specificity can be defined as the ability of the model to identify a structure from the unstable within energy changes, unstable within thermostability, affected activity, and disease potential.

An ideal ROC curve will approach the top left corner; therefore, a higher AUC means a better classifier. ROC curves are helpful for comparisons between different classifiers since they consider all thresholds. Eq (2) presents a logistic regression model used to predict protein structures (4IBY, 1TSR) using the variables in AUTOMUTE 2.0.

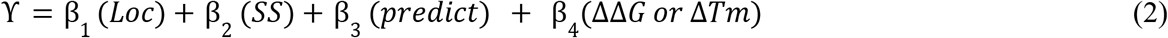

Y is a logit categorical binary variable representing 1TSR/4IBY structures. B_1_ (Loc) is a categorical variable referring to the depth of the residue in the protein structure (surface is referred to as S, the subsurface is referred to as U, and buried is referred to as B). B_2_(SS) a categorical variable referred to as secondary structure assessment for the residue (Beta-sheet is referred to as S, alpha-helix referred to as H, coil is referred to as C). B_3_(predict) categorical variable referred to the prediction status for the residue. B_4_(ΔΔG or ΔTm) is applied only ΔΔG (kcal/mol) for stability change, and ΔTm (°C) is for thermostability change. When diagonal, the ROC curve means the predictor variables (SC, TC, AC, PD) do not predict the structures. In other words, the model’s predictability is the same as a coin toss with AUC = 0.5.

In this paper, we created a silica mutagenesis file with 3515 entries for the DNA binding domain for P53. Then, we run the data with the following four classifiers in AUTO-MUTE: stability change (SC), thermostability change (TC), activity change (AC), and disease potential (DP). This study examined the prediction outcome of two structures for the P53 protein. Statistical methods such as ROC and UAC can be used by comparing 1TSR and 4IBY comprehensive structure prediction from AUTO-MUTE 2.0 software.

## RESULT

The paper presents various measurements to score the performance of the AUTO-MUTE. Four different predictors were used to predict the two structures. Numeric values were presented by stability change and thermostability change, and the classified values were presented by activity and disease potential. Therefore, it is necessary to set thresholds for stability change and thermostability change to classify the prediction. A classification threshold is set at −1 kcal/mol for stability change (ΔΔG), according to Jia et al. (2015). Values greater than −1 are considered stable, while values less than −1 are considered unstable. The thermostability-change threshold value (ΔTm) is set to −3°C (Jia, Yarlagadda, and Reed 2015). A value greater than −3°C is classified as “stable,” while a value less than −3 is classified as “unstable.”

The confusion matrix for AUTO-MUTE 2.0 is shown in Table 1. The calculation of the True-positives (TP), True-negatives (TN), False-positives (FP), and False-negatives (FN) between 1TSR and 4IBY. 84.5% of data in the SC dataset are (TP) and (TN) between 1TSR and 4IBY. However, only 15.5% of (FP) and (FN) are between the two structures. The thermostability presents 85.3% of the prediction (TP + TN) and 14.7% of the prediction (FP+FN). Also, 85.3% of the prediction is classified the same (TP+TN) for the Activity change, and 14.7% is differently classified (FP+FN). Moreover, the disease potential is around 73.1% of the prediction data is classified the same (TP+TN), and 26.9% is differently classified (FP+FN). Generally, it has been observed that 88% to 73% of the prediction is ideally classified, and less than 26% of predicted datasets are differently classified between the two P53 PDB structures.

**Table 1.** Confusion matrices for the four predictors using the AUTO-MUTE 2.0 program

| Stability-Change (SC) |  |  |  |  |
| --- | --- | --- | --- | --- |
|  | Stable<br>(4IBY) | % | Unstable<br>(4IBY) | % |
| Stable (1TSR) | 1414 | 40.2% | 271 | 7.7% |
| Unstable<br>(1TSR) | 275 | 7.8% | 1555 | 44.2% |
| Total | 3515 |  |  |  |
| Thermostability-Change (TC) |  |  |  |  |
|  | Stable<br>(4IBY) | % | Unstable<br>(4IBY) | % |
| Stable (1TSR) | 1468 | 41.8% | 232 | 6.6% |
| Unstable<br>(1TSR) | 286 | 8.1% | 1529 | 43.5% |
| Total | 3515 |  |  |  |
| Activity-Change (AC) |  |  |  |  |
|  | Unaffected<br>(4IBY) | % | Affected<br>(4IBY) | % |
| Unaffected<br>(1TSR) | 2403 | 68.4% | 227 | 6.5% |
| Affected<br>(1TSR) | 290 | 8.3% | 595 | 16.9% |
| Total | 3515 |  |  |  |

| Disease-Potential (DP) |  |  |  |  |
| --- | --- | --- | --- | --- |
|  | Neutral<br>(4IBY) | % | Disease<br>(4IBY) | % |
| Neutral (1TSR) | 1297 | 36.9% | 418 | 11.9% |
| Disease (1TSR) | 528 | 15.0% | 1272 | 36.2% |
| Total | 3515 |  |  |  |

The (SC) and (TC) histograms are presented in figures (1 and 2). The graphs show the histogram of the distribution of residues for different structures for stable and unstable datasets. The distribution considers each residue based on certain criteria (secondary structure assessment and residue depth location). The statistical analysis of SC and TC histograms indicates that 1TSR and 4IBY structures do not differ significantly. The p-value of stability-change (p-value= 0.1919 for stable, p-value= 0.1505 for unstable) and thermostability (p-value= 0.3313 for stable, p-value= 0.0695 for unstable). The permutation test was conducted for stability-change (p-value= 0.1505 for stable, p-value= 0.1476 for unstable) and thermostability (p-value= 0.3327 for stable, p-value= 0.0963 for unstable) which also indicated that there is no significant difference between the two structures (table 2). Differences in resolution between the two structures did not impact the results of computational mutagenesis.

**Figure 1.**
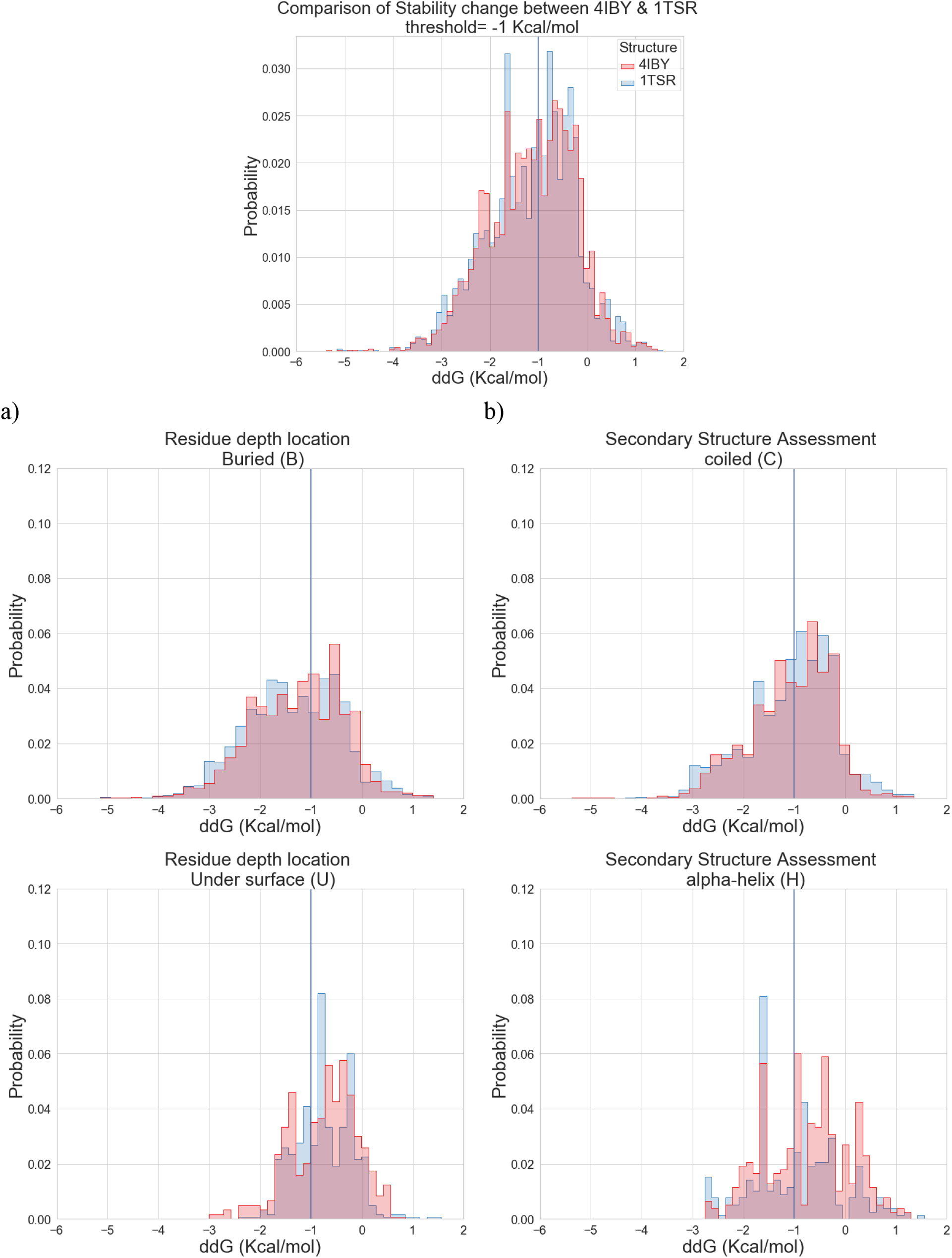
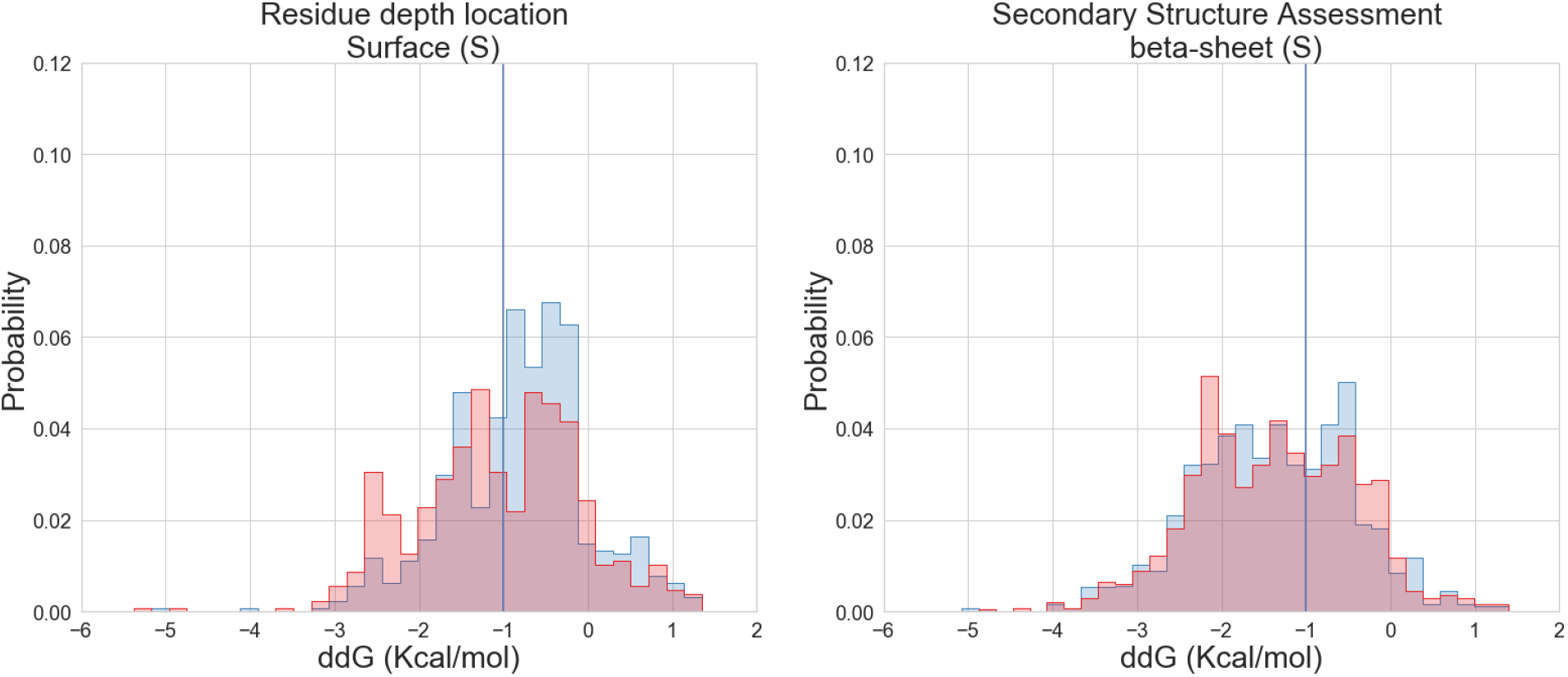
The histogram shows the distribution of stability change (ddG)for 4IBY and 1TSR with a threshold set to −1 kcal/mol, where less than this value is considered as unstable. The red color represents 4IBY, and the blue color represents 1TSR. The histograms show the locations of the residues on the protein structure where they occur in the stable dataset. Section a) contains three graphs that indicate the location of each residue site based on its classification as buried (B), under the surface (U), or on the surface (S). Three graphs in the b) section of the presentation show the distribution of the three types of secondary structure assessment (SSA), which were employed to classify the stable dataset as coiled (C), alpha-helix (H), or beta-sheet (S).

**Table 2.** Statistical comparison of stability changes and thermostability change for the t-test and permutation tests for stable and unstable

| Predictors | T-test (p-value) |  | Permutation test (p-value) |  |
| --- | --- | --- | --- | --- |
|  | Stable | Unstable | Stable | Unstable |
| SC | 0.1919 | 0.1505 | 0.1505 | 0.1476 |
| TC | 0.3316 | 0.0695 | 0.3327 | 0.0963 |

**Table 3.** Summary of the performance of the predictive models.

| Predictors | R-squared | Coeff. | Std err | P>\|t\| | Confident<br>(0.025) | Confident<br>(0.975) |
| --- | --- | --- | --- | --- | --- | --- |
| SC | 0.001246 |  |  |  |  |  |
| Location |  | -0.0151 | 0.023 | 0.502 | -0.059 | 0.029 |
| Secondary |  | -0.0457 | 0.022 | **0.035** | -0.088 | -0.003 |
| ddG |  | -0.1095 | 0.049 | **0.025** | -0.205 | -0.014 |
| Predict |  | -0.058 | 0.089 | 0.510 | -0.234 | 0.117 |
| TS | 0.002317 |  |  |  |  |  |
| Location |  | -0.0762 | 0.028 | **0.007** | -0.131 | -0.021 |
| Secondary |  | -0.0550 | 0.024 | **0.022** | -0.102 | -0.008 |
| dTm |  | -0.0354 | 0.009 | **0.000** | -0.053 | -0.018 |
| Predict |  | 0.1417 | 0.065 | **0.029** | 0.015 | 0.269 |
| AC | 0.001151 |  |  |  |  |  |
| Location |  | 0.0127 | 0.026 | 0.625 | -0.038 | 0.064 |
| Secondary |  | -0.0576 | 0.028 | **0.042** | -0.113 | 0.002 |
| Predict |  | 0.1583 | 0.063 | **0.011** | 0.038 | 0.281 |
| DP | 0.001226 |  |  |  |  |  |
| Location |  | -0.0479 | 0.021 | **0.020** | -0.088 | -0.008 |
| Secondary |  | 0.0181 | 0.022 | 0.406 | -0.025 | 0.061 |
| Predict |  | 0.1119 | 0.051 | 0.105 | 0.012 | 0.211 |

The predictive models all have low standard errors, indicating that the means are compact and accurately represent the true population mean. As indicated by the p-value of less than 0.05, this coefficient is statistically significant. According to the SC predictive model, secondary structure and ddG are not affected by structure prediction. The TC predictive model does not influence the secondary structure and location of ddG, but positively influences the predicted result. For the AC model, influence just predicts. Finally, the DP predictive model does not affect the location of the residue depth. The summary indicates that the predictive modes have little effect on the outcome. Therefore, the parameter from the predictive modes is independent of these variables. Thus, the prediction cannot predict the structure from the parameters. Although there are differences between the two structures, the result of the mutagenesis prediction does not differ in either structure.

Figure 6 presents the performance and ROC curves for each logistic regression model. The AUC values range between 0.47 and 50% of the data for AUTOMUTE predictive modes. As a result, AUCs demonstrate that the separation between the two structures cannot be determined using residue depth location, secondary structure assessment status, ddG (dTm). Furthermore, the similarity of the two structures is high even when using comprehensive mutagenesis prediction. Therefore, it is challenging to distinguish between the two structures based on the predicted parameters.

**Figure 3.**
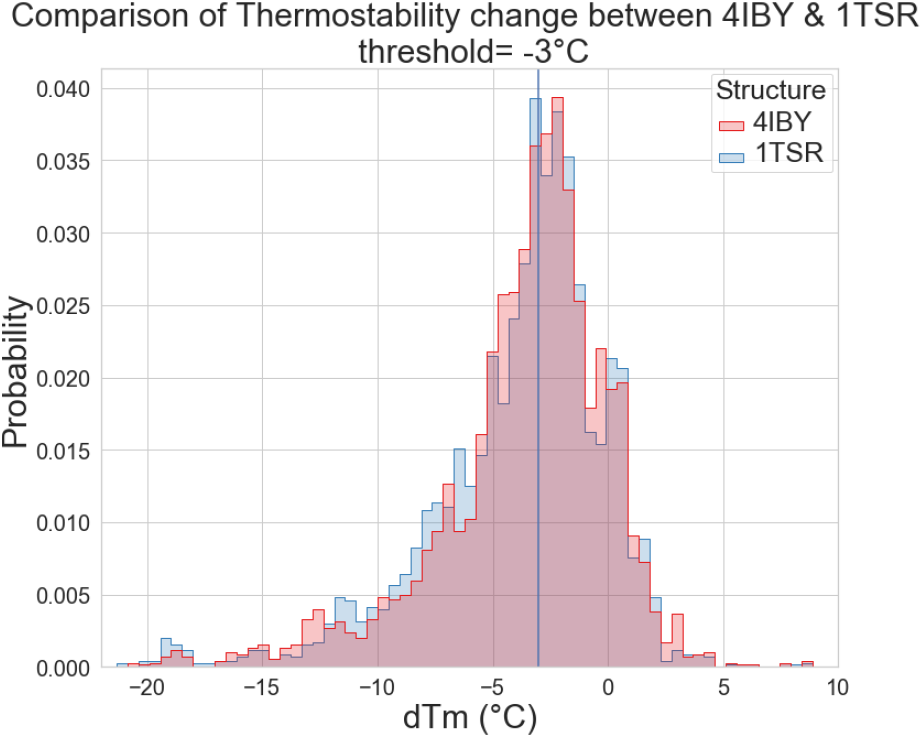
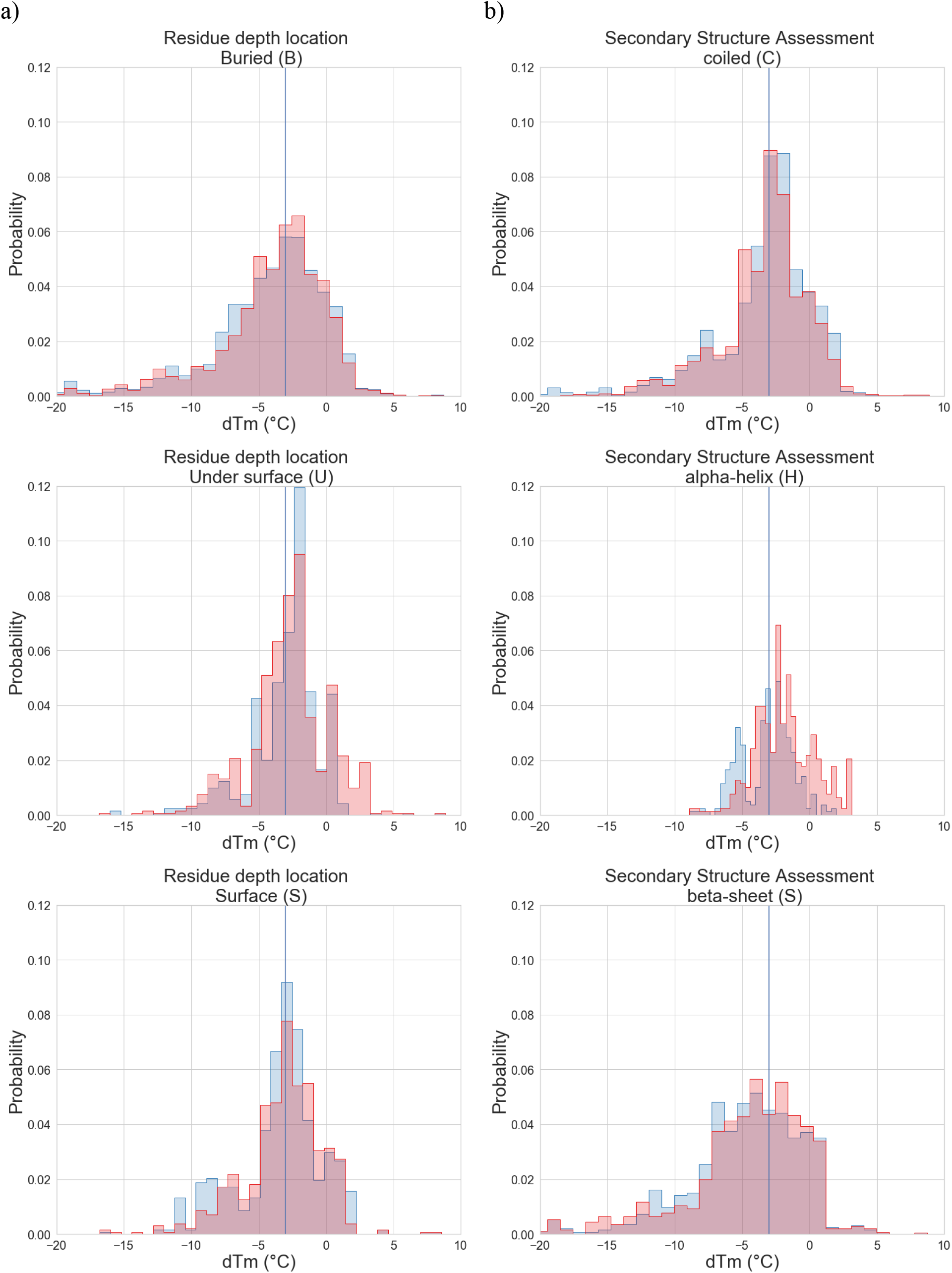
The histogram shows the distribution of thermostability (dTm) for 4IBY and 1TSR with a threshold set to −3°C, where less than this value is considered as unstable. The red color represents 4IBY, and the blue color represents 1TSR. The histograms show the locations of the residues on the protein structure where they occur in the stable dataset. Section a) contains three graphs that indicate the location of each residue site based on its classification as buried (B), under the surface (U), or on the surface (S). In addition, three graphs in the b) section of the presentation show the distribution of the three types of secondary structure assessment (SSA), which were employed to classify the stable dataset as coiled (C), alpha-helix (H), or beta-sheet (S).

**Figure 6.**
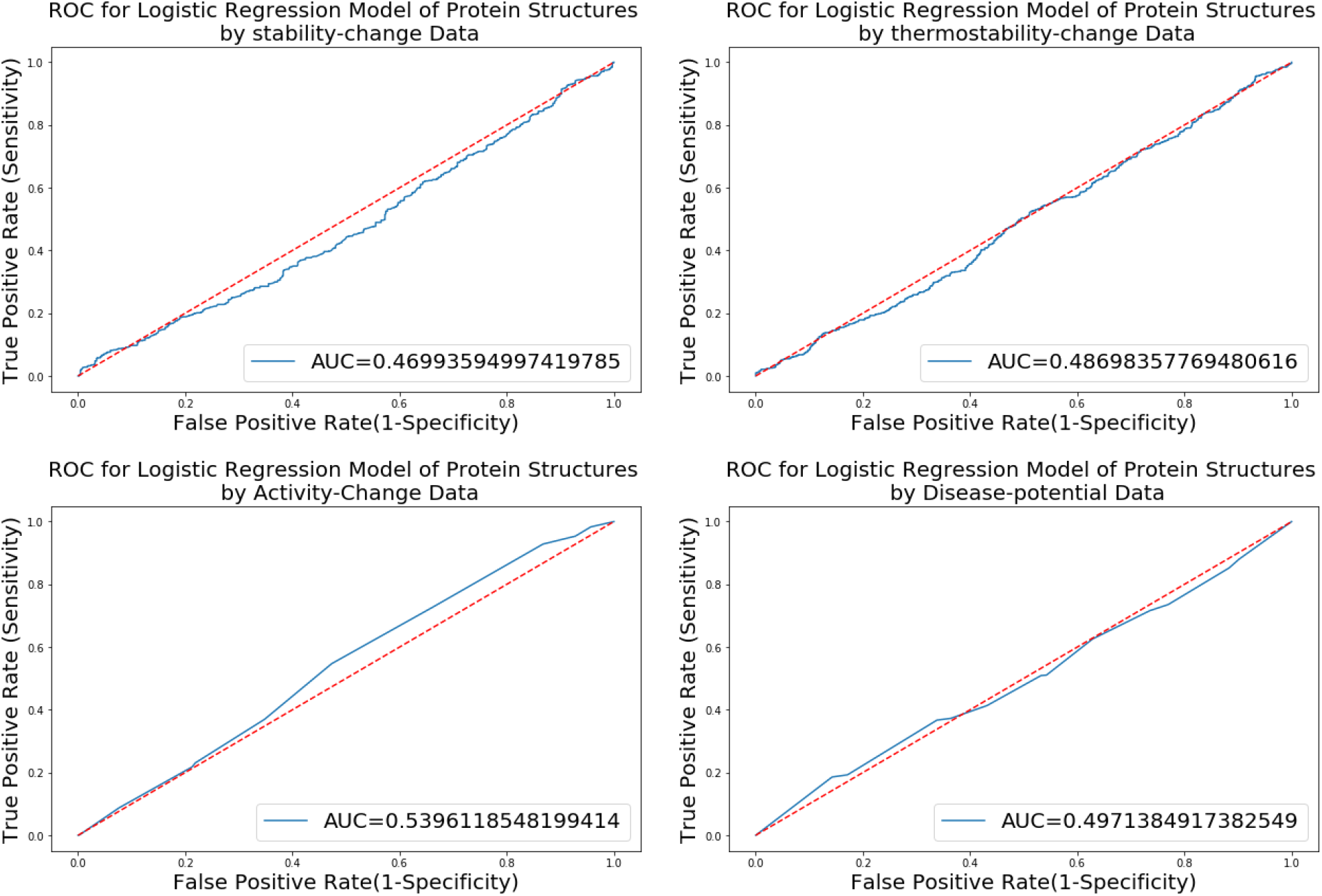
The ROC curve AUC values for each predictive model are calculated. The AUC for stability change is 0.47, thermostability change is 0.49, activity change is 0.54., and disease potential is 0.50

## DISCUSSION

Protein Data Bank (PBD) contains tens of thousands of protein structures. Several proteins in the database have more than one structural representation. Observed differences between the structures were caused by the experimental data metrics, particularly coverage and resolution. Nevertheless, the computation of mutagenesis in two different structures in resolution is satisfactory. We aimed to compare two crystal structures of the p53 protein with identical coverage but different resolutions in this article. 1TSR presents a resolution of 2.24Å, while 4IBY presents 1.4Å.

Here we discuss the results of the combined mutation of the two structure proteins 1TSR and 4IBY. Utilization of different predictive modes in AUTOMUTE, including stability changes (SC), thermostability changes (TC), activity changes (AC), and disease potentials (DP). The four predictive modes’ outcomes were examined using permutation analysis. Both structures showed 16 to 25% differences in the comprehensive mutagenesis prediction outcome, indicating both false positives and false negatives. Using the ROC curve assists in determining if these differences are significant. The results of the logistics regression were determined by using prediction variables to distinguish between the 1TSR and 4IBY structures. The AUC for the four classifiers was estimated to be 0.47-0.50. Thus, it is challenging to differentiate between the two structures based on the predicted parameters. As a result of the logistic regression, the two structures are so similar that they cannot be discriminated against.

There is no doubt that high resolution is a crucial aspect of x-ray crystallography to obtain the three-dimensional protein structure. The two protein structures present different levels of resolution. It is, however, evident from the statistical results that the difference is not a significant change. As a result of this study, it can be concluded that x-ray crystallography is an advanced and sensitive method, and the results of proteins at different resolutions produce similar results. Despite the minor difference in the structure of the protein resolution, studying low-resolution structures for any target protein could still be effective. Moreover, to validate this result, it would be beneficial to examine a large variety of datasets with different resolutions.

